# Evaluation and optimization of sequence-based gene regulatory deep learning models

**DOI:** 10.1101/2023.04.26.538471

**Authors:** Abdul Muntakim Rafi, Daria Nogina, Dmitry Penzar, Dohoon Lee, Danyeong Lee, Nayeon Kim, Sangyeup Kim, Dohyeon Kim, Yeojin Shin, Il-Youp Kwak, Georgy Meshcheryakov, Andrey Lando, Arsenii Zinkevich, Byeong-Chan Kim, Juhyun Lee, Taein Kang, Eeshit Dhaval Vaishnav, Payman Yadollahpour, Random Promoter DREAM Challenge Consortium, Sun Kim, Jake Albrecht, Aviv Regev, Wuming Gong, Ivan V. Kulakovskiy, Pablo Meyer, Carl de Boer

## Abstract

Neural networks have emerged as immensely powerful tools in predicting functional genomic regions, notably evidenced by recent successes in deciphering gene regulatory logic. However, a systematic evaluation of how model architectures and training strategies impact genomics model performance is lacking. To address this gap, we held a DREAM Challenge where competitors trained models on a dataset of millions of random promoter DNA sequences and corresponding expression levels, experimentally determined in yeast, to best capture the relationship between regulatory DNA and gene expression. For a robust evaluation of the models, we designed a comprehensive suite of benchmarks encompassing various sequence types. While some benchmarks produced similar results across the top-performing models, others differed substantially. All top-performing models used neural networks, but diverged in architectures and novel training strategies, tailored to genomics sequence data. To dissect how architectural and training choices impact performance, we developed the *Prix Fixe* framework to divide any given model into logically equivalent building blocks. We tested all possible combinations for the top three models and observed performance improvements for each. The DREAM Challenge models not only achieved state-of-the-art results on our comprehensive yeast dataset but also consistently surpassed existing benchmarks on *Drosophila* and human genomic datasets. Overall, we demonstrate that high-quality gold-standard genomics datasets can drive significant progress in model development.

## Introduction

In eukaryotes, transcription factors (TFs) play a crucial role in regulating gene expression and are critical components of the *cis*-regulatory mechanism (1–6). TFs compete with nucleosomes and each other for DNA binding and can enhance each other’s binding through biochemical cooperativity and mutual competition with nucleosomes (7–10). While the field has made substantial progress in characterizing regulatory mechanisms (11–19), a quantitative understanding of *cis*-regulation remains a major challenge. *Cis*-regulatory complexity grows with the square of the number of TFs involved due to the number of potential pairwise interactions between TFs (6). For example, since the yeast *Saccharomyces cerevisiae* has about 8.5x fewer TFs than humans (∼200 vs. ∼1,700) (1,20), its *cis*-regulatory code is theoretically about ∼72x less complex, making it an ideal system to test our understanding of eukaryotic *cis*-regulation.

Neural Networks (NNs) have shown immense potential in modeling and predicting gene regulation. While different network architectures, such as convolutional neural networks (CNNs) (11,12,14,19,21), recurrent neural networks (RNNs) (22), and transformers (15,17,18,23), have been used to create genomics models, there is limited research on how NN architectures and training strategies affect their performance for genomics applications. Standard datasets provide a common benchmark to evaluate and compare algorithms, leading to improved performance and continued progress in the field (24). For instance, the computer vision and natural language processing (NLP) fields have seen an ongoing improvement of NNs facilitated by gold-standard datasets, such as the ImageNet data (24), MS COCO (25), etc. In contrast, because genomics models are often created *ad hoc* for analyzing a specific dataset, it often remains unclear whether a model’s improved performance results from improved model architecture or better training data. In many cases, the models created are not directly comparable to previous models due to substantial differences in the underlying data used to train and test them.

To address the lack of standardized evaluation and continual improvement of genomics models, we organized the Random Promoter DREAM Challenge (26). Here, we asked the participants to design sequence-to-expression models and train them on expression measurements of promoters with random DNA sequences. The models would receive regulatory DNA sequence as input and use it to predict the corresponding gene expression value. We designed a separate set of sequences to test the limits of the models and provide insight into model performance. The top-performing solutions in the challenge exceeded performance of all previous state-of-the-art models for similar data. Our evaluation across various benchmarks revealed that, for some sequence types, model performances are approaching the previously-estimated inter-replicate experimental reproducibility for this datatype (13), while considerable improvement remains necessary for others. The top-performing models included features inspired by the nature of the experiment and the state-of-the-art models from computer vision and NLP, while incorporating novel training strategies that are better suited to genomics sequence data. To determine how individual design choices affect performance, we created a *Prix Fixe* framework that enables modular testing of individual model components, revealing further performance gains. Finally, we benchmarked the top-performing DREAM models on *Drosophila* and human datasets, including predicting expression and open chromatin from DNA sequence, where they consistently surpassed existing state-of-the-art performances. Recognizing the potential of these models to further the field, we are making all DREAM Challenge models available in an accessible format.

## Results

### The Random Promoter DREAM challenge and dataset

To generate the competition training data, we conducted a high-throughput experiment to measure the regulatory effect of millions of random DNA sequences (**Methods**). Prior research has shown that random DNA can display activity levels akin to genomic regulatory DNA, due to the incidental occurrence of numerous TF binding sites (13,23,27). Here, we cloned 80 bp random DNA sequences into a promoter-like context upstream of a yellow fluorescent protein (YFP), transformed the resulting library into yeast, grew the yeast in chardonnay grape must, and measured expression by fluorescent activated cell sorting (FACS) and sequencing, as previously described (13,28,29) (**Methods**). This resulted in a training dataset of 6,739,258 random promoter sequences and their corresponding mean expression values.

We provided these data to the competitors, who could use them to train their model, with two key restrictions. First, competitors were not allowed to use external datasets in any form to ensure that all models are trained on the same dataset. Second, ensemble predictions were also disallowed as they would almost certainly provide a boost in performance but without providing any insight into the best model types and training strategies.

We evaluated the models on a set of “test” sequences designed to probe the predictive ability of the models in different ways. The measured expression levels driven by these sequences were quantified in the same way as the training data but in a separate experiment with more cells sorted per sequence (∼100), yielding more accurate estimated expression levels compared to the training data measurements, and providing higher confidence in the challenge evaluation. The test set consisted of 71,103 sequences from several promoter sequence types. We included both random sequences and sequences from the yeast genome to get an estimate of performance difference between the random sequences in the training domain and naturally evolved sequences. We also included sequences designed to capture known limitations of previous models trained on similar data, namely sequences at the high and low expression extremes and sequences designed to maximize the disagreement between the predictions of a previously developed CNN and a physics-informed neural network (“biochemical model”) (13,23). We previously found that predicting changes in expression between closely related sequences (i.e., nearly identical DNA sequences) is substantially more challenging, and so we included subsets where models had to predict changes that result from single nucleotide variants (SNVs), perturbations of specific TF binding sites, and tiling of TF binding sites across background sequences (13,23). Each test subset was given a different weight when scoring the submissions, proportional to the number of sequences in the set and how important we considered it to be (**Table 2**). For instance, predicting the effects of SNVs on gene expression is a critical challenge for the field due to its relevance to complex trait genetics (30). Accordingly, a substantial number of SNV sequence pairs were included in the test set, and SNVs were given the highest weight. Within each sequence subset, we determined model performance using Pearson *r*^2^ and Spearman ρ, which captured the linear correlation and monotonic relationship between the predicted and measured expression levels (or expression differences), respectively. The weighted sum of each performance metric across test subsets yielded our two final performance measurements, which we call *Pearson Score* and *Spearman Score*.

Our DREAM Challenge ran for 12 weeks in the summer of 2022 and included two evaluation stages: the public leaderboard phase and the private evaluation phase (**Fig. 1A**). The leaderboard opened six weeks into the competition and allowed teams to submit up to 20 predictions on the test data per week. At this stage, we used 13% of the test data for leaderboard evaluation and displayed only the overall Pearson *r*^2^, Spearman ρ, *Pearson Score,* and *Spearman Score* to the participants, while keeping the performance on the promoter subsets and the specific sequences used for the evaluation hidden. The participating teams achieved increasing performance each week (**Supplementary Fig. 1**), showcasing the effectiveness of such challenges in motivating the development of better machine learning models. Over 110 teams across the globe competed in this stage. At the end of the challenge, 28 teams submitted their models for final evaluation. We used the rest of the test data (∼87%) for the final evaluation (**Fig. 1B,C; Supplementary Fig. 2**).

**Figure 1:**
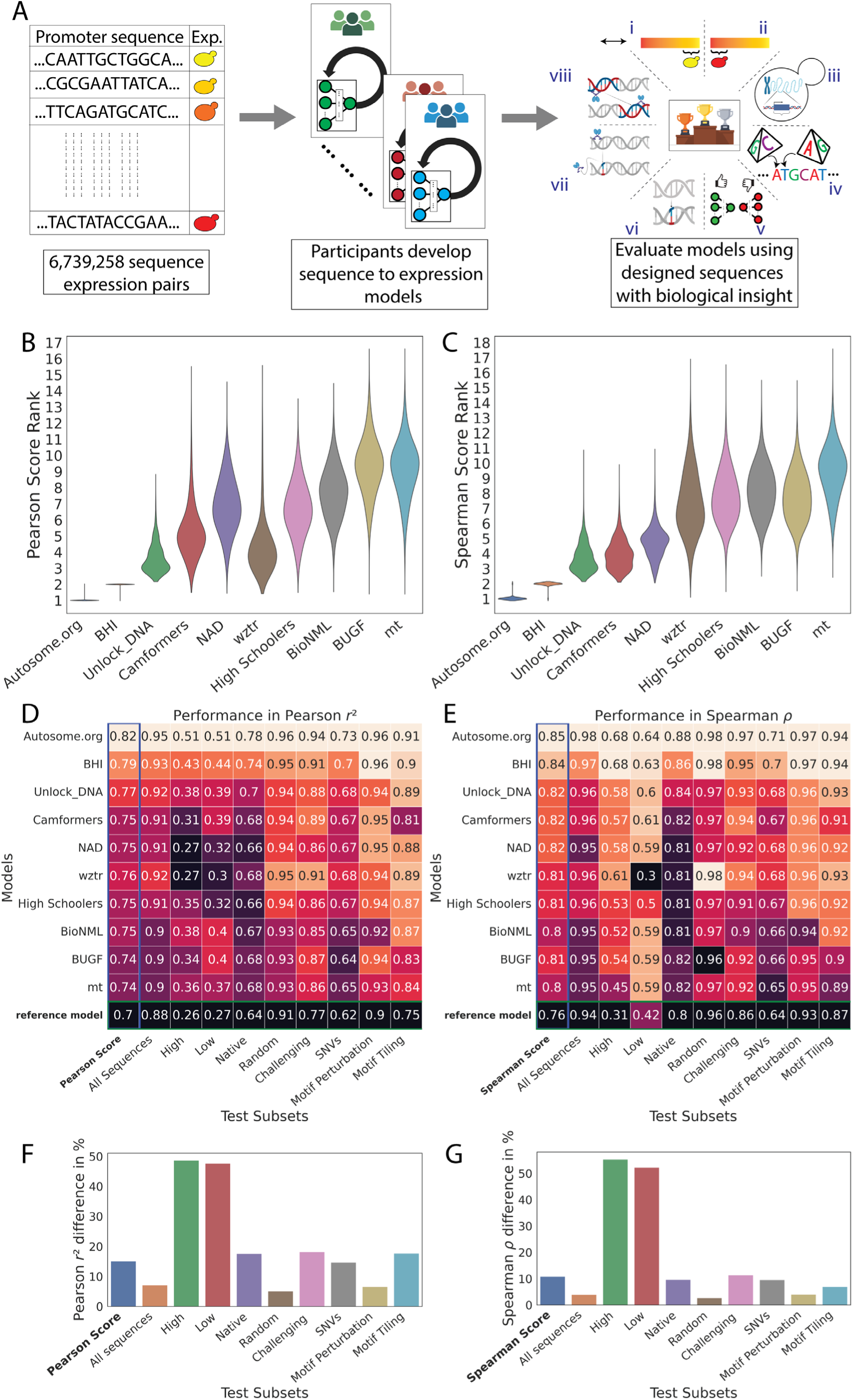
Overview of the challenge. (**A**) Competitors received a training dataset of random promoters and corresponding expression values (left). They continually refined their models and competed for dominance in a public leaderboard (middle). At the end of the challenge, they submitted a final model for evaluation (right) using a test dataset consisting of eight sequence types: (i) high expression, (ii) low expression, (iii) native, (iv) random, (v) challenging, (vi) SNVs, (vii) motif perturbation, and (viii) motif tiling. (**B, C**) Bootstrapping provides a robust comparison of the model predictions. Distribution of ranks in *n*=10,000 samples from the test dataset (*y*-axes) for the top-performing teams (*x*-axes), for (B) Pearson Score and (C) Spearman Score. (**D,E**) Performance of the top-performing teams in each test data subset. Model performance (colour and numerical values) of each team (*y*-axes) in each test subset (*x*-axes), for (D) Pearson *r*^2^ and(E) Spearman ⍴. Heatmap colour palettes are min-max normalized column-wise. (**F,G**) Performance disparities observed between the best and worst models (*x*-axes) in different test subsets (*y*-axes) for (F) Pearson *r*^2^ and (G) Spearman ⍴. The calculation of the percentage difference is relative to the best model performance for each test subset.

### Innovative model designs surpass the state-of-the-art

We retrained the transformer model architecture of Vaishnav et al. (12), the previous best-performing model for this type of data, on the challenge data and used as a reference in the leaderboard (“reference model”). The overall performance of top submissions, all NNs, was substantially better than the reference model. Despite recent prominence of attention-based architectures (23), only one of the top five submissions in the challenge used transformers, placing 3^rd^. The best-performing submissions were dominated by fully convolutional NNs, with 1^st^, 4^th^, and 5^th^ places taken by them. The best-performing solution was based on the EfficientNetV2 architecture (31,32), and the 4^th^ and 5^th^ solutions were based on the ResNet architecture (33). Moreover, all teams used convolutional layers as the starting point in their model design. A recurrent neural network (RNN) with bidirectional long-short-term memory (Bi-LSTM) layers (34,35) placed 2^nd^. While the teams broadly converged on many similar training strategies (e.g., using Adam (36) or AdamW (37) optimizers), they also had substantial differences (**Table 1**).

**Table 1:** Breakdown of the top-performing models into key components.

| Participant team name | NN architecture type | Input encoding and channels | Input flanking region length | Usage of reverse strand during model training | Train validation split | Parameters (millions) | Optimizer | Loss function | Learning Rate Scheduler | Metric |
| --- | --- | --- | --- | --- | --- | --- | --- | --- | --- | --- |
| Autosome.org | CNN (EfficientNetV2 (32)) OHE | OHE [6:bases/NC*/RC*] | 70 | Data aug. (additional channel) + Model (additional channel) | 100-0 | 1.9 | AdamW (37) | Kullback-Leibler divergence | One Cycle LR | $r, \rho^\#$ |
| BHI | CNN + RNN (Bi-LSTM) (34) | OHE [4:bases] | 30 | Post-hoc conjoined setting (41) | 100-0 | 6.8 | AdamW (37) | Huber | Cosine Anneal LR | $r, \rho^\#$ |
| Unlock_DNA | Transformer | OHE [6:bases/N*/M*] | 20 | Input to model (concat. with forward strand) | 95-5 | 47.4 | Adam (36) | MSE + custom | One Cycle LR | $r$ |
| Camformers | CNN (ResNet (33)) | OHE [4:bases] | 30 | None | 90-10 | 16.6 | AdamW (37) | L1 | Reduce LR On Plateau | $r, \rho$ |
| NAD | CNN + Transformer | GloVe (38) [128] | 0 | None | 90-10 | 15.5 | AdamW (37) + GSAM (42) | smooth L1 | Linear LR | $r$ |
| wztr | CNN (ResNet (33)) | OHE [4:bases] | 62 | Input to model (concat. with forward strand) | 99-1 | 4.8 | Adam (36) | MSE | Reduce LR On Plateau | $r$ |
| High Schoolers Are All You Need (High Schoolers) | CNN + Transformer + MLP | OHE [4:bases] | 31 | Model (RC parameter sharing) (41) | 98-2 | 4.7 | Adam (36) + SWA (43) | MSE | Multi Step LR | $r$ |
| BioNML | Vision Transformer (44) | OHE [4:bases] | 30 | Model (RC parameter sharing) (41) | 86-14 | 78.7 | Adamax (36) + L2 regularizer | Huber | Multi Step LR | $r, CoD$ |
| BUGF | Transformer | OHE [6:bases/N*/P*] | 32 | None | 94-6 | 4.5 | RAdam (45) | Multi-label focal loss (46) + custom | None | $r$ |
| mt | GRU (47) | OHE | 62 | Model (RC | 99.8-0.2 | 3.1 | Adam (36) | binary cross. | None | $r,$ |
|  | +CNN | [6:bases/<br>N*/P*] |  | parameter sharing) (41) |  |  |  |  |  | CoD<sup>#</sup> |

\*NC: If the sequence was present in more than one cell, 0 for all bases, otherwise 1; RC: If the sequence is reverse-complemented, 1 for all bases, otherwise 0; N: If a base is unknown, 1 for that base, otherwise 0; P: If a base has been padded to maintain fixed input length, 1 for that base, otherwise 0; M: If a base is masked 1 for that base otherwise 0.
<sup>#</sup>These teams employed the metrics in a cross-validation setting to determine the optimal number of epochs for their models and ultimately saved the model weights after running for the $n$ epochs, without relying on validation metric scores. In contrast, other teams utilized validation metric scores to select the best-performing model.

The competing teams introduced several innovative approaches to solve the expression prediction problem. Autosome.org, the best-performing team, transformed the task into a soft-classification problem by training their network to predict a vector of expression bin probabilities, which was then averaged to yield an estimated expression level, effectively recreating how the data were generated in the experiment. They also used a distinct data encoding method by adding channels to the traditional 4-channel one-hot encoding (OHE) of the DNA sequence used by most teams. The two additional channels indicated (i) whether the sequence provided as input was likely measured in only one cell (which results in an integer expression value), and (ii) whether the input sequence is being provided in the reverse complement orientation. Furthermore, Autosome.org’s model, with only two million parameters, challenged the current trend of designing deep NNs with increasingly numerous parameters, demonstrating that efficient design considerably reduces the necessary number of parameters. Autosome.org and BHI were distinct in training their final model on the entirety of the provided training data (i.e., no sequences withheld for validation) for a prespecified number of epochs (determined previously using cross-validation using validation subsets). Unlock_DNA, the 3^rd^ team, took a novel approach by randomly masking 5% of the input DNA sequence and having the model predict both the masked nucleotides and gene expression. This approach used the masked nucleotide predictions as a regularizer, adding a reconstruction loss to the model loss function, which stabilized the training of their large NN. BUGF, the 9^th^ team, used a somewhat similar strategy where they randomly mutated 15% of the sequence and calculated an additional binary cross-entropy loss predicting whether any bp in the sequence had been mutated. The 5^th^ team, NAD, employed GloVe (38) to generate embedding vectors for each base position and used these vectors as inputs for their NN, whereas the other teams utilized traditional one-hot encoded DNA sequences. Two teams, SYSU-SAIL-2022 (11^th^) and Davuluri lab (16^th^), attempted to train DNA language models (39) on the challenge data, by pretraining a BERT language model (40) on the challenge data and subsequently using the BERT embeddings to train an expression predictor.

### Test sequence subsets reveal model disparities

Analysis of model performance on the different test subsets revealed distinct and shared challenges for the different models. The top two models were ranked 1^st^ and 2^nd^ (sometimes with ties) for each test subset regardless of score metric, showcasing that their superior performance cannot be attributed to any single test subset (**Fig. 1D,E**). Further, the rankings within each test subset sometimes differed between Pearson Score and Spearman Score, reinforcing that these two measures capture performance in distinct ways (**Fig. 1D,E**).

While the ranking of models was similar for both random and native sequences, the differences in model performance were greater for native yeast sequences than random sequences. Specifically, performance differed between models by as much as 17.6% for native sequences, but only 5% for random sequences (Pearson *r*^2^, **Fig. 1F**; Similarly 9.6% (native) *vs*. 2.7% (random) for Spearman ⍴; **Fig. 1G**). This suggests that top models have learned more of the regulatory grammar that evolution has produced. Further, the substantial discrepancy between performance on native and random sequences suggests that there is yet more regulatory logic to learn (although the native DNA have lower sequence coverage, presumably due to their higher repeat content, likely reducing quality data; **Supplementary Fig. 3**).

Models were also highly variable in their ability to accurately predict variation within the extremes of gene expression. The cell sorter has a reduced signal-to-noise ratio at the lowest expression levels, and the sorting bin placement can truncate the tails of the expression distribution (6,12). Overall, model performance was most variable across teams in these subsets, suggesting that the challenge models were able to overcome these issues to varying degrees. For example, the median difference in Pearson *r*^2^ between highest and lowest performance was ∼48% for high and low test subsets and 16% for the others (**Fig. 1F,G**).

The models also varied in their ability to predict expression differences between closely related sequences (**Fig. 1D,E**; “SNVs”, **Supplementary Fig. 4**; **Supplementary Fig. 5**), with more substantial differences in model performance for subtler changes. Specifically, the % difference between best and worst in Pearson *r*^2^ and Spearman ⍴ were respectively, 6.5% and 4% for Motif Perturbation, 17.7% and 7% for Motif Tiling, and 14.6% and 9.6% for SNVs, suggesting that the top-performing models better captured the subtleties of cis-regulation. This is consistent with our understanding of the subtlety of the impact: perturbing TFBSs (“Motif Peturbations” where we mutate sequences strongly matching the cognate motif for an important TF or vary the number of binding sites) is a comparatively large perturbation and could be predicted with simple models that capture the binding of these TFs and can count TFBS instances. However, when TFBSs are tiled across a background sequence, the same TFBS is present in every sequence, and the model must have learned how its position affects its activity, as well as capturing all the secondary TFBSs that are created or destroyed as the motif is tiled (13). Finally, SNVs are even harder to predict because nearly everything about the sequence is identical but for a single nucleotide that may affect the binding of multiple TFs in potentially subtle ways.

### *Prix Fixe* framework reveals optimal model configurations

The top three solutions from the DREAM Challenge were distinguished both by their substantial improvement in performance compared to other models and their distinct approaches to data handling, preprocessing, loss calculations, and diverse neural network layers—encompassing convolutional, recurrent, and self-attention mechanisms. To identify the factors underlying their performances, we developed a *Prix Fixe* framework that breaks down each solution into distinct modules and, by selecting one of each module type, tests arbitrary combinations of the modules from each solution (**Fig. 2A**). We re-implemented the top three solutions within this framework and found that 45 out of 81 possible combinations were compatible. We removed specific test-time processing steps unique to each solution that were not comparable across solutions. Finally, we retrained all compatible combinations using the same training and validation data, addressing the issue that some original solutions had used the entire dataset for training. Our approach facilitated a systematic and fair comparison of the individual contributions of different components to overall performance.

**Figure 2:**
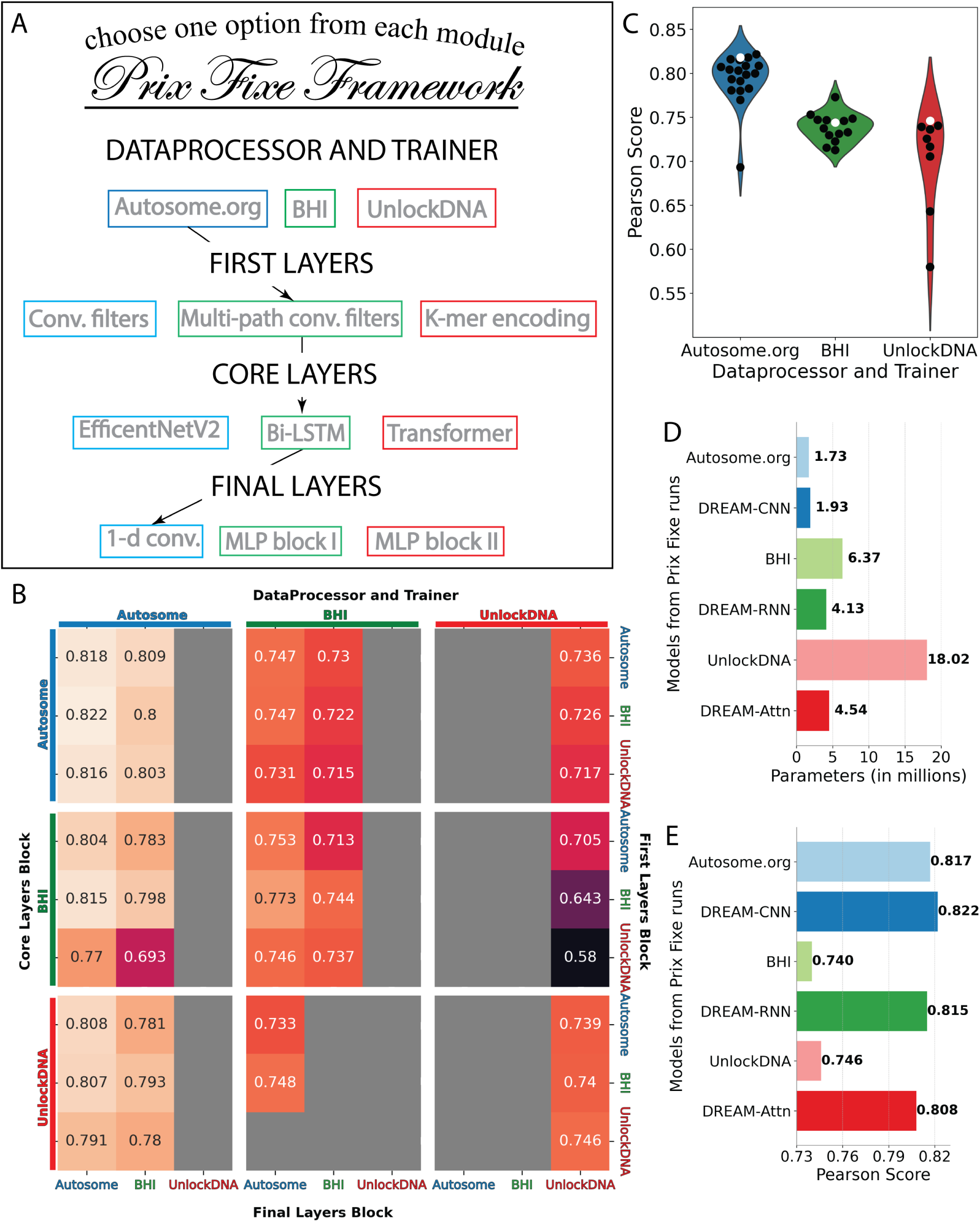
Dissecting the optimal model configurations through a *Prix Fixe* framework. **(A)** The framework deconstructs each team’s solution into modules, enabling modules from different solutions to be combined. (**B**) Performance in Pearson Score from the *Prix Fixe* runs for all combinations of modules from the top three DREAM Challenge solutions. Each cell represents the performance obtained from a unique combination of Core Layers Block (major rows, left), DataProcessor and Trainer (major columns, top), First Layers Block (minor rows, right), and Final Layers Block (minor columns, bottom) modules. Grey cells denote combinations that were either incompatible or did not converge during training. **(C)** Performance (Pearson Score; *y*-axis) of the three DataProcessor and Trainer modules (*x*-axis and colours) for each *Prix Fixe* model including the respective module (individual points). Original model combinations are indicated by white points, while all other combinations are in black. **(D)** Number of parameters (*x*-axis) for the top three DREAM Challenge models-Autosome.org, BHI, and UnlockDNA-along with their best-performing counterparts (based on Core Layers Block): DREAM-CNN, DREAM-RNN, and DREAM-Attn in the *Prix Fixe* runs (*y*-axis). **(E)** As in D, but showing each model’s Pearson Score (*x*-axis).

Our analysis revealed both the source of Autosome.org’s exceptional performance as well as the interplay of different model components and their potential for further optimization. ’The BHI and UnlockDNA NNs saw significant improvement in performance when retrained using Autosome.org’s soft-classification paradigm (**Fig. 2B,C; Supplementary Fig. 6; Supplementary Fig. 7**). Moreover, each team’s model architecture could be optimized further, resulting in models that achieved better performance (**Fig. 2C**) using the same core blocks but with similar or fewer parameters (**Fig. 2D**). However, except for Autosome.org’s DataProcessor and Trainer module, no other module component dominated the others and their performance appeared to depend on what other modules they were combined with (**Supplementary Fig. 8**). For each of Autosome.org, BHI, and UnlockDNA’s core blocks, we named the optimal *Prix Fixe* model as DREAM-CNN, DREAM-RNN, and DREAM-Attn, respectively.

### Optimized models outperform state-of-the-art for other species and data types

To determine whether the model architectures and training strategies we optimized on yeast data would generalize to other species, we next applied them to *Drosophila melanogaster* and human datasets on a diverse set of tasks. First, we tested their ability to predict gene regulatory activity measured in *D. melagnogaster* (in the context of a developmental and a housekeeping promoter) in a STARR-seq MPRA. This fundamentally represents the same sequence-to-expression problem the models were designed to solve, despite the different organism (*Drosophila* vs. yeast), experimental measurement approach (RNA-seq vs. cell sorting), longer sequence (249 bp vs. 150 bp), smaller datasets (∼500k thousand vs. 6.7 million), and the transition from a single-task to a multi-task framework (two promoter types). We compared the DREAM-optimized models to DeepSTARR (48), a state-of-the-art CNN model based on the Basset (21) architecture and specially developed for predicting the data we are using in this benchmark (UMI-STARR-seq (49) in *D. melanogaster* S2 cells (48,50)). For a robust comparison, we trained the models using cross validation and always evaluated on the same held-out test data (**Methods**). Our models consistently outperformed DeepSTARR across both developmental and housekeeping transcriptional programs (**Fig. 3A**), with the DREAM-RNN model surpassing the performance of DREAM-CNN and DREAM-Attn.

**Figure 3:**
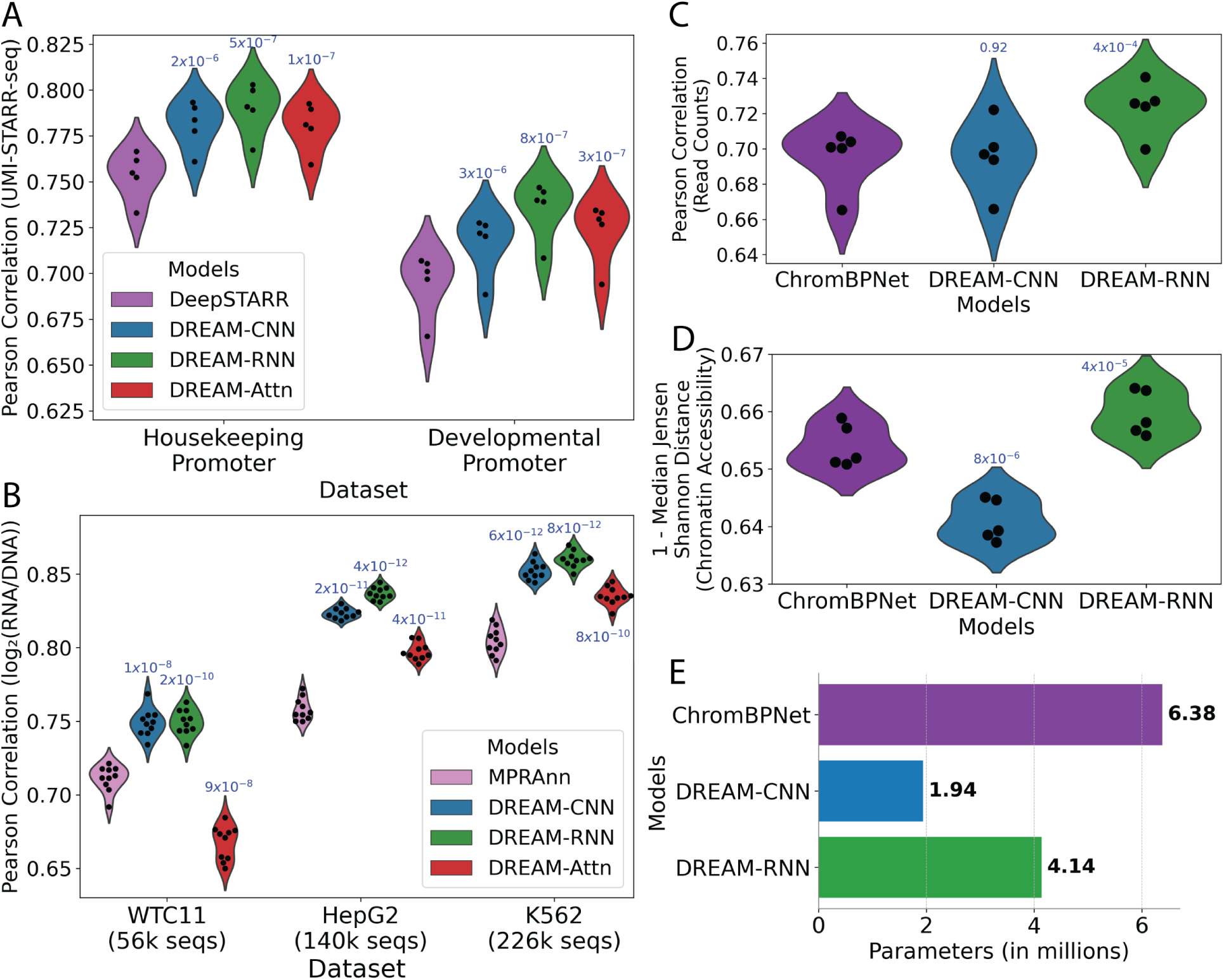
DREAM Challenge models beat existing benchmarks on *Drosophila* and human datasets. **(A)** *D. melanogaster* STARR-seq (48) prediction. Pearson correlation for predicted *vs*. actual enhancer activity for held out data (*y*-axis) for two different transcriptional programs (*x*-axis) for each model (colours). **(B)** Human MPRA (51) prediction. Pearson correlation for predicted *vs*. actual expression for held out data (*y*-axis) for MPRA datasets from three distinct human cell types (*x*-axis) for each model (colours). **(C, D)** Human accessibility (bulk K562 ATAC-seq) (52,53) prediction. For each model (x-axis and colours), model performance (*y*-axes) in shown in terms of both **(C)** Pearson correlation for predicted *vs*. actual read counts per element and **(D)** 1-median Jensen-Shannon distance for predicted *vs*. actual chromatin accessibility profiles across each element. In **A-D**, points represent folds of cross validation, performance is evaluated on held-out test data, and p-values representing t-tests (paired, two-sided) comparing the previous state-of-the-art model to the optimized models are shown atop model performance distributions. **(E)** Comparison of number of parameters (*x*-axis) for different models used in chromatin accessibility prediction task.

To further validate the generalizability of our models, we next trained the DREAM-optimized models on lentivirus-based massively parallel reporter assays (lentiMPRAs) that tested *cis* regulatory elements (CREs) across three human cell types: hepatocytes (HepG2), lymphoblasts (K562), and induced pluripotent stem cells (iPSCs; WTC11) (51). Here, our models had to capture more complex regulatory activity from vastly smaller datasets (∼56-226 thousand *vs.* 6.7 million). We compared the models against MPRAnn (51), a CNN model optimized for these specific datasets (**Methods**). All the models were trained using cross validation and evaluated on held-out test data in the same way MPRAnn was originally trained (51). The DREAM-optimized models substantially outperformed MPRAnn, with the performance difference widening with more training data (**Fig. 3B**). The only exception was DREAM-Attn, which did not outperform MPRAnn on the smallest dataset (WTC11; 56k sequences). Again, DREAM-RNN demonstrated the best performance among our models, especially for larger datasets.

To evaluate the models on a distinct prediction task that still relates to CRE function, we evaluated our optimized models on the task of predicting open chromatin. Specifically, we compared our optimized models to ChromBPNet (53–55), a BPNet-based (16) model that predicts ATAC-seq signals across open chromatin regions. Here, the input DNA sequences were ∼14 times longer than the yeast promoters on which the DREAM models were optimized (2,114 vs 150 bp), and the models were now tasked with simultaneously predicting the overall accessibility (read counts) and accessibility profile (read distribution) for a central 1,000 bp section, rather than predicting a single expression value. While DREAM-Attn could not be trained because the memory requirement for the attention block became too large with such a long input sequence, we trained and evaluated the other DREAM-optimized models and ChromBPNet on K562 bulk ATAC-seq data (52) (**Methods**). DREAM-RNN outperformed ChromBPNet substantially in both read counts prediction and chromatin accessibility predictions (**Fig. 3C,D**), highlighting the adaptability of our models even on significantly different cis-regulatory data types. DREAM-CNN, on the other hand, performed on par with ChromBPNet (53) in read count predictions (**Fig. 3C**) but was less effective in predicting chromatin accessibility profiles (**Fig. 3D**).

Notably, the architectures and training paradigms of the DREAM-optimized models were changed minimally for these evaluations, and the only modifications made were required for the prediction head (e.g. because the model outputs differed) or to adapt to the smaller number of training sequences compared to the DREAM dataset (reducing the batch size and/or maximum learning rate; **Methods**). Importantly, DREAM-RNN outperformed the other *Prix Fixe* optimized models in all of these secondary benchmarks (**Fig. 3A-D**), highlighting its excellent generalizability.

## Discussion

The random promoter DREAM Challenge 2022 presented a unique opportunity for participants to propose novel model architectures and training strategies for modeling regulatory sequences. The participants trained sequence-to-expression models on millions of random regulatory DNA sequences and their corresponding expression measurements. A separate set of designed sequences were used to evaluate these models and test their limits. Remarkably, 19 models from the DREAM Challenge outperformed the previous state-of-the-art (23) (**Supplementary Fig. 4**), with the majority employing unique architectures and training strategies. To systematically analyze how model design choices impact their performance, we developed the *Prix Fixe* framework, where models were abstracted to modular parts, enabling us to combine modules from different submissions to identify the key contributors to model performance. We applied the *Prix Fixe* framework to the top three models from the challenge that varied significantly in their NN architectures (CNN, RNN, self-attention) and training strategies, and were able to construct improved models in each case.

The training strategies for NNs had as significant an impact on model performance as the network architectures themselves (**Fig. 2C; Supplementary Fig. 8**). In the *Prix Fixe* runs, training the network to predict expression as distributions using soft-classification rather than as precise values helped models capture more of cis-regulation. These findings argue for a balanced focus not only on network architectures but also on the optimization of training procedures and redefinition of prediction tasks.

Notably, the top-performing models from the DREAM Challenge demonstrated that simpler neural network architectures with fewer parameters, if optimized well, can effectively capture most of cis-regulation, challenging the current trend towards increasingly complex NNs. For instance, three of the top five submissions did not use transformers, including the best-performing team (which also had the fewest parameters of the top 10). Also, by employing our *Prix Fixe* framework, we successfully designed models that not only consisted of similar or fewer parameters but also achieved superior performance compared to their counterparts (**Fig. 2D,E**). Further, these DREAM-optimized models consistently outperformed previous state-of-the-art models on other cis-regulatory tasks, despite having comparable (and often fewer) parameters (**Fig. 1D**, **Fig. 3E**, **Supplementary Fig. 9**). These results indicate that in genomics applications, building better models does not necessarily require increased complexity, but rather depends on efficient and effective optimization.

In the DREAM challenge, we observed varied results across test subsets that illustrate the complexity in evaluating cis-regulatory models effectively. For instance, performance on random sequences, which are in the same domain as the training data (also random sequences), was relatively uniform (**Fig. 1D,E**). Conversely, shifting the domain to native sequences highlights the disparities between models, as the relative frequencies of various regulatory mechanisms likely differ, a consequence of their evolutionary origin (**Fig. 1D,E**). This indicates that a model that excels in modeling overall cis-regulation may still perform poorly for sequences involving certain regulatory mechanisms that are difficult to learn from the training data, leading to incorrect predictions of biochemical mechanisms and variant effects for sequences that use these mechanisms. This emphasizes the importance of multifaceted evaluation of genomics models (56), and designing specific datasets that test the limits of these models.

To continually improve genomics models, there is a need for standardized, robust benchmarking datasets. The DREAM Challenge dataset addresses this need and the impact such standardized datasets can have is demonstrated by the DREAM-optimized models’ generalizability across different *Drosophila* and human datasets and tasks without additional model tuning. Our dataset accompanied by the *Prix Fixe* framework stand as valuable resources for the continued exploration and development of innovative neural network architectures and training methodologies specifically crafted for DNA sequences. Further, the modular nature and proven generalizability of the DREAM-optimized models will enable other researchers to easily apply them to other genomics problems.

## Data availability

Data is available at NCBI’s GEO: GSE254493.

## Code availability

Open source code for our models is available at https://github.com/de-Boer-Lab/random-promoter-dream-challenge-2022

## Methods

### Designing the test sequences

High- and low-expression sequences were designed as described in Vaishnav et al. (23). Native test subset sequences were designed by sectioning native yeast promoters into 80 bp fragments (13). Random sequences were sampled from a previous experiment where the tested DNA was synthesized randomly (as in the training data) and quantified (13). Challenging sequences were designed by maximizing the difference between the expressions predicted by a convolutional neural network model (23) and a biochemical model (a type of physics-informed neural network) (13); these sequences represented the pareto front of the differences in expression between models when optimizing populations of 100 sequences at a time for 100 generations using a genetic algorithm with a per-base mutation rate of 0.02 and recombination rate of 0.5 using DEAP (57) and a custom script (github.com/de-Boer-Lab/CRM2.0/blob/master/GASeqDesign.py). Most of the SNVs represent sequence trajectories from Vaishnav et al. (23), but also include random mutations added to random, designed, and native promoter sequences. Motif Perturbation included Reb1 and Hsf1 perturbations. Sequences with perturbed Reb1 binding sites were created by inserting Reb1 consensus binding sites (strong or medium affinity; sense and reverse complement orientations), and then adding between 1 and 3 SNVs to each possible location of each motif occurrence and inserting canonical and mutated motif occurrence into 10 randomly generated sequences at position 20/80. Sequences with Hsf1 motif occurrence were designed by tiling random background sequences with between 1 and 10 Hsf1 monomeric consensus sites (ATGGAACA), added sequentially from both the right and left of the random starting sequences, or added individually within each of the possible 8 positions, or similarly tiling/inserting between 1-5 trimeric Hsf1 consensus sites (TTCTAGAANNTTCT). The Motif Tiling test subset sequences were designed by embedding a single consensus for each motif (poly-A: AAAAA, Skn7: GTCTGGCCC, Mga1: TTCT, Ume6: AGCCGCC, Mot3: GCAGGCACG, and Azf1: TAAAAGAAA) at every possible position (with the motif contained completely within the 80-bp variable region) and orientation for three background sequences as described in de Boer et al. (13).

### Quantifying promoter expression

High complexity random DNA libraries that comprised the training data were created as described previously (13), with the following modifications. The random promoter library in E. coli theoretically contained about 74 million random promoters and was transformed into S288c (delta URA3) yeast yielding 200 million transformants, which were selected in SD-Ura media. 1 L of chardonnay grape must (filtered) was inoculated with the pool to an initiate OD600 of 0.05 and grown at room temperature without continual shaking, with the culture diluted as needed with fresh chardonnay grape must to maintain OD below 0.4, for a total growth time of 48 hours and having undergone >5 generations. Prior to each OD, the culture was gently agitated to decarbonate it, waiting for the resulting foam to die down before agitating again, and continuing until no more bubbles were released. Yeast were then sorted and sequencing libraries prepared as described previously (13), with sequencing libraries pooled and sequenced on an Illumina NextSeq.

Data processing for both N80 (training) and designed (test) libraries were done as described before (13), except that the expression levels of the designed library sequences were estimated using MAUDE (58), using the read abundance in each sorting bin as input, and estimating the initial abundance of each sequence as the average relative abundance of that sequence across all bins.

### Competition rules

1. Only the provided training data could be used to train models. Models had to train from scratch without any pre-training on external datasets to avoid overfitting to sequences present in the test data (e.g., some sequences in the test data are derived from extant yeast promoters).
2. Reproducibility was a prerequisite for all submissions. The participants had to provide the code and instructions to reproduce their models. We retrained the top-performing solutions to validate their performance.
3. Augmenting the provided training data was allowed. Pseudo-labeling the provided test data was not allowed. Using the test data for any purpose during training was not allowed.
4. Ensembles were not allowed.

For detailed information on the competition and its guidelines, please visit the DREAM Challenge webpage (https://www.synapse.org/#!Synapse:syn28469146/wiki/617075).

### Performance evaluation metric

We calculated the Pearson *r^2^* and Spearman ⍴ between predictions and measurements for each sequence subset. Pearson *r* captures the linear correlation between predictions and measured expression levels while being robust to the scaling differences that occur between training and test sequences; the training and the test data are not on the same scale but are linearly related by, *y* = *mx* + *b*, where *m* and *b* are constants that are affected by sorting bin placement during the experiment. We also used Spearman ⍴, which captures the monotonic relation between predictions and measured expression levels while being robust to outliers. The weighted sum of each performance metric across promoter types yielded our two final performance measurements, which we call *Pearson Score* and *Spearman Score*.

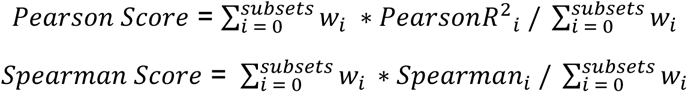

Here, *w*_*i*_ is the weight used for the *i*-th test subset (**Table 2**). *PearsonR*^2^_*i*_ and *Spearman*_*i*_ are, respectively, the square of Pearson coefficient and the Spearman coefficient for sequences in *i*-th subset.

**Table 2:** Summary of the test subsets.

| Subset | no. of sequences | Weight in evaluation metric | Description |
| --- | --- | --- | --- |
| All sequences | 71103 | 1 | All sequences in the test data. |
| High | 968 | 0.3 | Sequences designed to have high expression. |
| Low | 997 | 0.3 | Sequences designed to have low expression. |
| Native | 997 | 0.3 | Sequences that are present in the yeast genome. |
| Random | 6349 | 0.3 | Random DNA sequences. |
| Challenging | 1953 | 0.5 | Sequences designed to maximize the differences between a convolutional model and a biochemical model trained on the same data. |
| SNVs | 44340 pairs | 1.25 | Two sequences that differ by only a single base. |
| Motif Perturbation (Reb1+Hsf1) | 3287 pairs | 0.3 | Two sequences that differ due to perturbations to specific known TF binding site. |
| Motif tiling | 2624 pairs | 0.4 | Two sequences that differ due to tiling known TF binding sites across random sequences. |

### Bootstrapping analysis of model performance

In order to determine the relative performance of the models, we performed a bootstrapping analysis. Here, we sampled 10% of the test data 10,000 times and, for each sample, calculated the performance of each model and the rankings of the models for both Pearson and Spearman Scores. We averaged the ranks from both metrics to decide their final ranks. Teams Autosome.org and BHI robustly outperformed the others, coming in 1^st^ and 2^nd^ place, respectively, with Autosome.org coming second to BHI in only 0.1% and 2.25% of the time for *Pearson Score* and *Spearman Score*, respectively. In none of the bootstraps did another team outperform Autosome.org or BHI. Interestingly, Pearson Score and Spearman Score do not agree after the top two teams, indicating that they capture performance in distinct ways. For instance, Unlock_DNA, the 3^rd^-place team, substantially outperformed 4^th^-place Camformers by *Pearson Score,* but performed slightly worse than Camformers by *Spearman Score*. We consider 5^th^-place to be a tie between NAD and wztr as their mean ranks (5.81105 and 5.8152, respectively) were very close.

### Description of the approaches used by the participants

In this section, we present an overview of the approaches employed by the participants in the challenge. For the top-performing teams, we provide a detailed description of their methodologies, while for the remaining teams, we offer a concise overview without repeating any details that have already been discussed for other teams. The performance of each approach is illustrated in **Supplementary Fig. 4** for easier comparison.

#### Autosome.org

The team reformulated the initial regression task as a soft-classification problem by replacing the initial target expression value with a vector of 18 probabilities corresponding to individual bins, assuming that they can be deduced from the normal distribution with mean and variance equal to (expression + 0.5) and 0.5, respectively. To obtain a predicted expression value for a sequence during the validation step, the predicted probabilities were multiplied by their bin numbers. They used one-hot encoding for the promoter sequences, where they added a separate binary channel explicitly marking the objects with integer (and thus likely imprecise) expression measurements. They also augmented the dataset with the reverse complementary sequences and added a separate binary channel to denote the supplied strand explicitly (forward or reverse complementary). The team also highlighted the impact of the training regime on the final performance, specifically, the advantage of using the OneCycleLR scheduler coupled with the AdamW optimizer. The proposed model was based on a fully-convolutional network inspired by EfficientNetV2 (32). The following architectural choices were used in the final model: (i) grouped convolution (59) instead of the depthwise convolution of the original EfficientNetV2, (ii) the standard residual blocks were substituted with residual channel-wise concatenations, (iii) a bilinear layer was inserted in the middle of the EfficientNetV2 SE-block. A detailed study on this architecture is presented in (60).

#### BHI

Their approach adopts a “sandwich” architecture consisting of a one-dimensional convolutional layer, a bidirectional long-short-term memory (Bi-LSTM) layer, and another convolutional layer. Each convolutional layer used different kernel sizes. Besides the model architecture, the team found that training details specialized for DNA sequence-based deep learning models were highly important for the overall performance. Among them, the most crucial was to use a ‘post-hoc conjoined’ setting (41), which imposes a reverse-complement equivariance to the model. Test-time augmentation was also effective. Predictions were made for an original sequence, its four shifted variants (generated by -2bp, -1bp, +1bp, and +2bp shifting), and their reverse-complement sequences, then those 10 predictions were averaged to make a final prediction. While training sequences over 110bp were trimmed to the right, sequences shorter than 110bp were randomly padded with the original vector sequences on both sides. This informative padding gave a nonnegligible performance boost. Finally, to be as unbiased as possible for the distribution of the test set, predictions were quantile-transformed using the distribution of expression levels in training data as post-processing.

#### Unlock_DNA

The team used an end-to-end Macaron-like Transformer encoder architecture with two half-step feed-forward (FFN) layers at the beginning and end of each encoder block. A separable 1D convolution layer was inserted after the first FFN layer and in front of the multi-head attention layer. The sliding k-mers from one-hot encoded sequences were mapped onto a continuous embedding, combined with the learned positional embedding and strand embedding (forward strand vs. reverse complement strand) as the sequence input. Along with the sequence input, several positions (32 in the final model) of "pseudo" expression values were added as the input, where all input expression values were zeros. The model predicted one expression value for each "pseudo" expression position and used the mean of the prediction of all positions as the final predicted expression value. A detailed study on this solution is presented on (61).

#### Camformers

This team used a CNN with residual connections. The model included six convolutional layers with three residual connections allowing the model to bypass every other layer. After the penultimate convolutional layer, a max pooling operation was added to reduce the model size and improve generalization. The output of the final convolutional layer was flattened and fed into a block of two dense layers, followed by a final dense layer outputting the predicted expression level. All layers except the last used a rectified linear unit activation.

#### NAD

The approach has two stages: (i) generating the embedding vectors for each base position using GloVe (38) and (ii) using the embedding vectors as input of neural networks to predict the gene expression level. The proposed model combines a convolutional neural network for feature extraction and a transformer for prediction.

#### WZTR

The team used a fully CNN-based architecture. The model begins with two convolutional layers, and six convolution blocks follow these layers. Each convolution block is constructed of 3 convolutional layers and an average pooling at the end. Each convolutional layer consists of a hybrid convolution (62), batch normalization, ReLU activation, and residual connection. Each hybrid convolution takes in a list of dilation values [1,2,4,6], with 4 convolutions processing the input in parallel. Finally, there are three fully connected layers and an output layer. A linear combination of 256 features extracted from all the previous operations on the sequence is used to generate the predicted expression.

#### High Schoolers Are All You Need (High Schoolers)

This team used a mix of CNN and transformer architectures, where the CNN was based on Residualbind’s (63) design, with a convolutional layer (with exponential activations) followed by a residual block comprised of a series of dilated convolutional layers with increasing dilation rates. It was followed by attention pooling, a transformer layer with relative positional encodings, and a standard MLP block.

#### BioNML

The underlying neural network was configured to have a relatively larger set of convolutional kernels and extra dictionaries of short k-mers for spotting potential enriched DNA sequence patterns. Strand-specific streams of these patterns were normalized and consolidated with Swish activation’s (64) fully learnable thresholding. The encoded patterns were fed into a ViT (44) like block but with transformer decoder type of connections and SwiGLU (65) activations for modeling any sequential interdependence. A set of suppressed signals of the encoded sequence-based patterns as queries for the transformer decoder blocks to respond to.

#### BUGF

A transformer model was used to predict the expression bin classes, as opposed to treating the problem as a regression problem. Random mutations were added to the sequence as an data augmentation strategy and the model was trained to predict where the mutations had been made to the input sequence. An auxiliary loss was calculated based on this prediction, which helped reduce overfitting.

#### mt

The approach uses GRU and CNNs to regress the strength of the targeted promoters using information encoded in the forward and reverse DNA strands.

#### SYSU-SAIL-2022

The team first trained a 3-layer BERT (40) using the top 20% of sequences in terms of expression. Then, the BERT embedding was used to train an expression predictor.

#### Wen Group

A deep neural network that adopted concepts of U-Net (66), Transformer, and Squeeze & Excitation blocks (67) was trained from end to end without any data augmentation.

#### Yuanfang Guan

A neural network that consisted of LSTM layers followed by attention layers was used to predict expression.

#### Metformin-121

A neural network based on bidirectional GRU was used to predict gene expression.

#### NGT4

A neural network based on XceptionNet (68) was used to predict gene expression. During training, the expression values were transformed evenly in the range of *i* – 0.5 < *x* < *i* + 0.5 (here, *i* is the integer expression) maintaining the ranking of sequences that was produced by a trained model.

#### Davuluri lab

The team utilized a transformer-based representation model named DNABERT (39) for predicting gene expression.

*DNABERT was pretrained on human genome, which violated the competition rules. However, we consider this to be an important benchmark that shows the limitation of DNA language models.

#### Wan&Barton_BBK

The team designed a model based on Temporal Convolutional Networks (69) to predict expression.

#### Peppa

The team designed a model based on the Enformer (15) that took 110 bp as input (compared to 200 kbp in the original) and included only 2M parameters (compared to ∼200M parameters of the original).

#### The Dream Team

A neural network was used that incorporated convolutional, multihead attention, and LSTM layers. During training, the integer expression values were transformed by replacing them with Normal(i,0.3) distribution, where i represents the expression.

#### Noisy-Chardonnay

A model composed of convolutional layers followed by BiLSTM layers was trained without any data augmentation to predict expression levels.

#### KircherLab

The team trained a simple convolutional neural network with a GC correction step on the training data to help the model focus its decisions on motifs within the sequence rather than the general nucleotide composition.

#### MadLab

The model is composed of three building blocks, namely a convolutional network, a transformer and a recurrent network.

#### Auth

A simple hybrid architecture combining a convolutional layer and a BiLSTM layer followed by two fully connected layers was used to predict expression.

#### UTKbioinformatics

A neural network based on BERT was used to predict expression.

#### DrAshokAndFriends

An attention based ConvLSTM (70) model was used for prediction.

#### QUT_seq2exp

A sequence embedding model, dna2vec (71), was applied on the promoter sequences (in a running manner on short k-mers), which are then subsequently used as features for a transformer-based deep neural network model.

#### Zeta

A transformer model was used for predicting expression.

##### *Prix Fixe* Framework

The *Prix Fixe* framework, implemented in Pytorch, facilitates the design and training of NNs by modularizing the entire process, from data-preprocessing to prediction, enforcing specific formats for module inputs and outputs to allow integration of components from different approaches. The different modules in the *Prix fixe* framework are as follows:

#### DataProcessor and Trainer

This DataProcessor class is dedicated to transforming raw DNA sequence data into a usable format for subsequent NN training. The DataProcessor can produce an iterable object, delivering a dictionary containing, a feature matrix ’x’ (input to the NN) and a target vector ’y’ (expected output). Additional keys can be included to support extended functionalities. Moreover, the DataProcessor can provide essential parameters to initiate NN blocks, such as determining the number of channels in the first layer.

The Trainer class manages the training of the NN. It processes batches of data from the DataProcessor and feeds them into the NN. It computes auxiliary losses, if necessary, alongside the main losses from the NN, facilitating complex loss calculation during training.

#### Prix Fixe Net

This module embodies the entirety of the NN architecture:

(i) First Layers Block: These are the primordial layers of the network. They may include initial convolutional layers or facilitate specific encoding mechanisms like k-mer encoding for the input.

(ii) Core Layers Block: This represents the central architecture components, housing elements like residual connections, LSTM mechanisms, and self-attention. The modular construction of this block also allows for versatile combinations, such as stacking a residual CNN block with a self-attention block.

(iii) Final Layers Block: This phase narrows the latent space to produce the final prediction, using layers like pooling, flattening, and dense layers. It computes the prediction and outputs it alongside the loss.

For all three blocks, the standard input format is (batch, channels, seqLen). The first two blocks yield an output in a consistent format (batch, channels, seqLen), whereas the last block delivers the predicted expression values. Each block can propagate their own loss. The whole framework is implemented in PyTorch.

To ensure fair comparison across solutions in the Prix Fixe framework, we removed specific test-time processing steps that were unique to each solution. We divided the DREAM Challenge dataset into two segments, allocating 90% sequences for training and 10% for validation. Using these data, we retrained all combinations that were compatible within the framework. Out of the 81 potential combinations, we identified 45 as compatible, and 41 of these successfully converged during training. Due to GPU memory constraints, we adjusted the batch sizes for certain combinations.

### DREAM-optimized models from *Prix Fixe* runs

#### DataProcessor and Trainer

Promoter sequences were extended at the 5’ end using constant segments from the plasmids to standardize to a length of 150 bp. These sequences underwent one-hot encoding into four-dimensional vectors. ’Singleton’ promoters, observed only once across all bins, were categorized with integer expression estimates. Considering the potential variability in these singleton expression estimates, a binary ’is_singleton’ channel was incorporated, marked as 1 for singletons and 0 otherwise. To account for the diverse behavior of regulatory elements based on their strand orientation relative to transcription start sites, each sequence in the training set was provided in both its original and reverse complementary forms, identified using the ’is_reverse’ channel (0 for original, 1 for reverse complementary). Consequently, the input dimensions were set at (batch, 6, 150).

The model’s training utilized the AdamW optimizer, set with a weight_decay of 0.01. The maximum learning rate of 0.005 was chosen for most blocks, while a conservative rate of 0.001 was applied to attention blocks due to the inherent sensitivity of self-attention mechanisms to higher rates. This learning rate was scheduled by the One Cycle Learning Rate Policy (OneCycleLR) (72), which featured two phases and employs the cosine annealing strategy. Training data was segmented into batches of size 1024, with the entire training procedure spanning 80 epochs. Model performance and selection were based on the highest Pearson r value observed in the validation dataset.

During prediction, the data processing mirrored the DataProcessor apart from setting ’is_singleton’ to 0. Predictions for both the original and reverse complementary sequences were then averaged.

##### *Prix Fixe* Net

###### (i) DREAM-CNN

First Layers Block: One-hot encoded input is processed through a 1D-CNN. Drawing inspiration from DeepFam (73), convolutional layers with kernel sizes of 9 and 15 are used, which mirror common motif lengths as identified by ProSampler (74). Each layer has a channel size of 256, uses ReLU activation, and incorporates a dropout rate of 0.2. The outputs of the two layers are concatenated along the channel dimension.

Core Layers Block: This segment contains six convolution blocks whose structure is influenced by the EfficientNet architecture. It contains modifications like replacing depthwise convolution with grouped convolution, using Squeeze and Excitation (SE) blocks (67), and adopting channel-wise concatenation for residual connections. The channel configuration starts with 256 channels for the initial block, followed by 128, 128, 64, 64, 64, and 64 channels (60).

Final Layers Block: The final block consists of a single point-wise convolutional layer followed by channel-wise Global Average Pooling and SoftMax activation.

###### (ii) DREAM-RNN

First Layers Block: same as DREAM-CNN.

Core Layers Block: The core utilizes a Bi-LSTM, designed to capture motif dependencies. The LSTM’s hidden states have dimensions of 320 each, resulting in 640 dimensions after concatenation. A subsequent CNN block, similar to the First Layer block, is incorporated.

Final Layers Block: same as DREAM-CNN.

###### (iii) DREAM-Attn

First Layers Block: It is a standard convolution with kernel size 7, followed by BatchNorm (75) and SiLU activation (76).

Core Layers Block: This block uses the Proformer (61), a Macaron-like Transformer encoder architecture, which employs a two half-step feed forward network (FFN) layers at the start and end of the encoder block. Additionally, a separable 1D convolution layer is integrated after the initial FFN layer and prior to the multi-head attention layer.

Final Layers Block: same as DREAM-CNN and DREAM-RNN.

### Human MPRA models

Within each of the three large-scale MPRA libraries, every sequence and its corresponding reverse complement are grouped together and these pairs were then distributed into ten distinct cross-validation folds to ensure both the forward and reverse sequences resided within the same fold. DREAM-CNN, DREAM-RNN, DREAM-Attn, and MPRAnn were trained using nine out of these ten folds, reserving one fold to evaluate the model’s performance. For every held-out test fold, nine models were trained, with one fold being dedicated for validation purposes while the remaining eight acted as training folds. Subsequent predictions from these nine models were aggregated, with the average being used as the final prediction for the held-out test data.

The training specifics of MPRAnn were followed as described in (51). For DREAM-CNN, DREAM-RNN, and DREAM-Attn, components that could not accommodate Agarwal et al. data were discarded. For instance, the ‘is_singleton’ channel is not relevant for MPRA data, loss calculation was done using mean squared error in place of KL divergence due to the infeasibility of transitioning the problem to soft classification. MPRAnn used a much smaller batch size than our DREAM-optimized Trainer (32 vs 1024), and so we reduced it to be the same as MPRAnn. No other alterations were made to either the model’s structure or the training paradigm.

### Drosophila UMI-STARRSeq models

The training specifics (model architecture and trainer) of DeepSTARR were followed as described (48). For DREAM-CNN, DREAM-RNN, and DREAM-Attn we used the exact setting as our runs in human MPRA datasets.

Only the five largest *Drosophila* chromosomes (Chr2L, Chr2R, Chr3L, Chr3R, ChrX) were used as test data. For every held-out test chromosome, the rest of the sequences were distributed into ten distinct train-validation folds, and DREAM-CNN, DREAM-RNN, DREAM-Attn, and DeepSTARR models (ten of each) were trained. Subsequent predictions from these ten models were aggregated, with the average being used as the final prediction for the held-out test chromosome.

### Human Chromatin accessibility models

We used five separate train-validation-test splits as proposed in (53) for ATAC-seq experiments on the human cell line K562 (52). For each of these partitions, we first trained five different bias models, one per fold, which are designed to capture enzyme-driven biases present in ATAC-seq profiles. Subsequently, ChromBPNet, DREAM-CNN, and DREAM-RNN models were trained for each fold, using the same bias models. For DREAM-CNN and DREAM-RNN, the prediction head from ChromBPNet (1-d convolution, cropping layer, average pooling layer, and a dense layer) was used in the Final Layers Block to make accessibility profile and read counts prediction. We modified the DREAM-optimized Trainer in this case to use the same batch size as ChromBPNet (from 1024 to 64), and a reduced maximum learning rate (from 5 x 10^-4^ to 5 x 10^-3^). No other alterations were made to either the model’s structure or the training paradigm.

For this task, we reimplemented DREAM-CNN and DREAM-RNN architectures in Tensorflow to ensure all models have same bias models. This methodological choice came at the cost of having to leave some components (input encoding, AdamW optimizer, etc.) out of the DREAM-optimized DataProcessor and Trainer. However, it ensured uniformity across models, leading to an unbiased comparison across the different architectures.

## Supplementary Figures

**Supplementary Figure 1:**
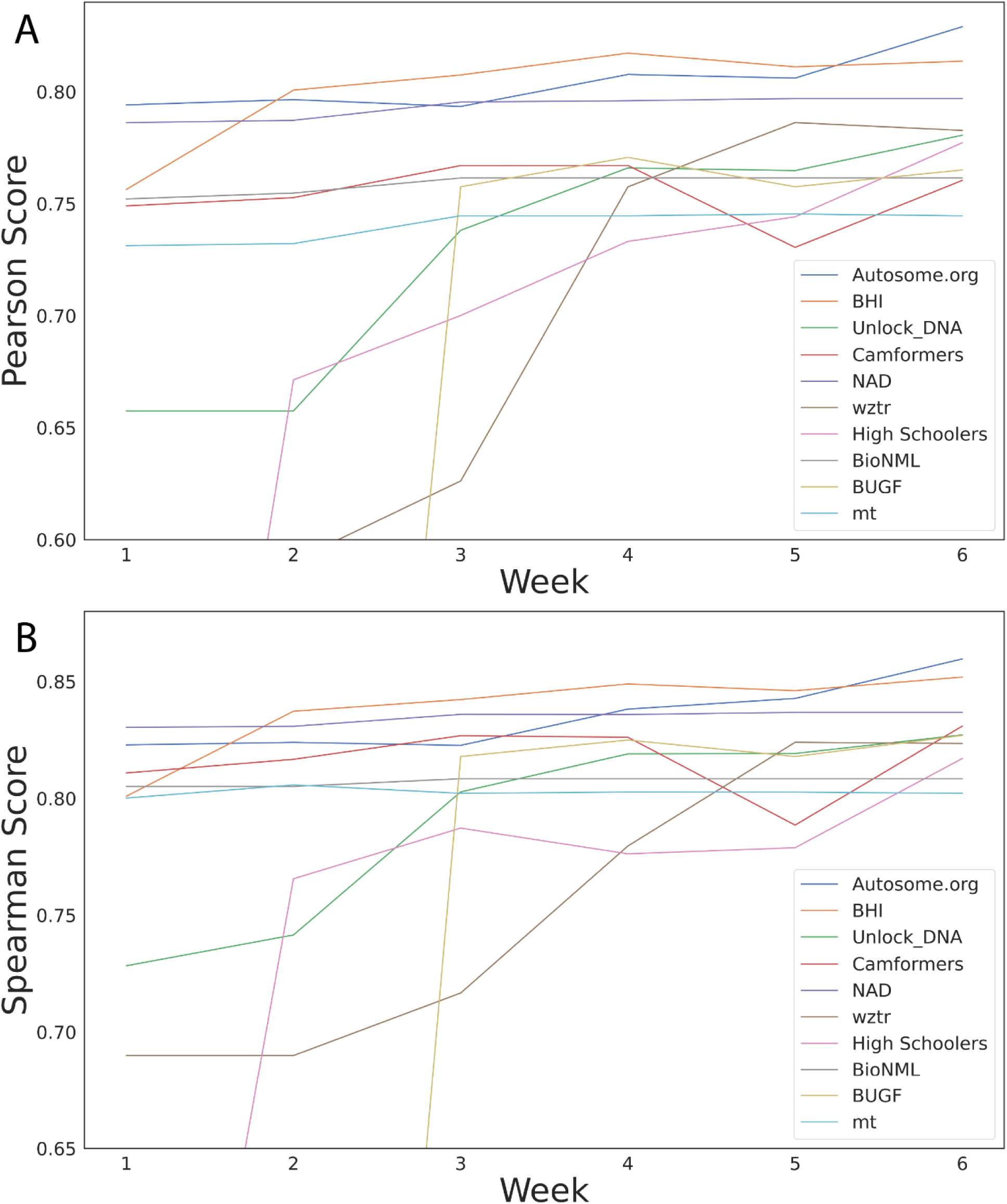
Progression of the top-performing teams’ performance in the DREAM Challenge public leaderboard. **(A,B)** Performance (*y*-axes) in (**A**) Pearson Score and (**B**) Spearman Score achieved by the participating teams (colours) each week (*x*-axes), showcasing the effectiveness of such challenges in motivating the development of better machine learning models.

**Supplementary Figure 2:**
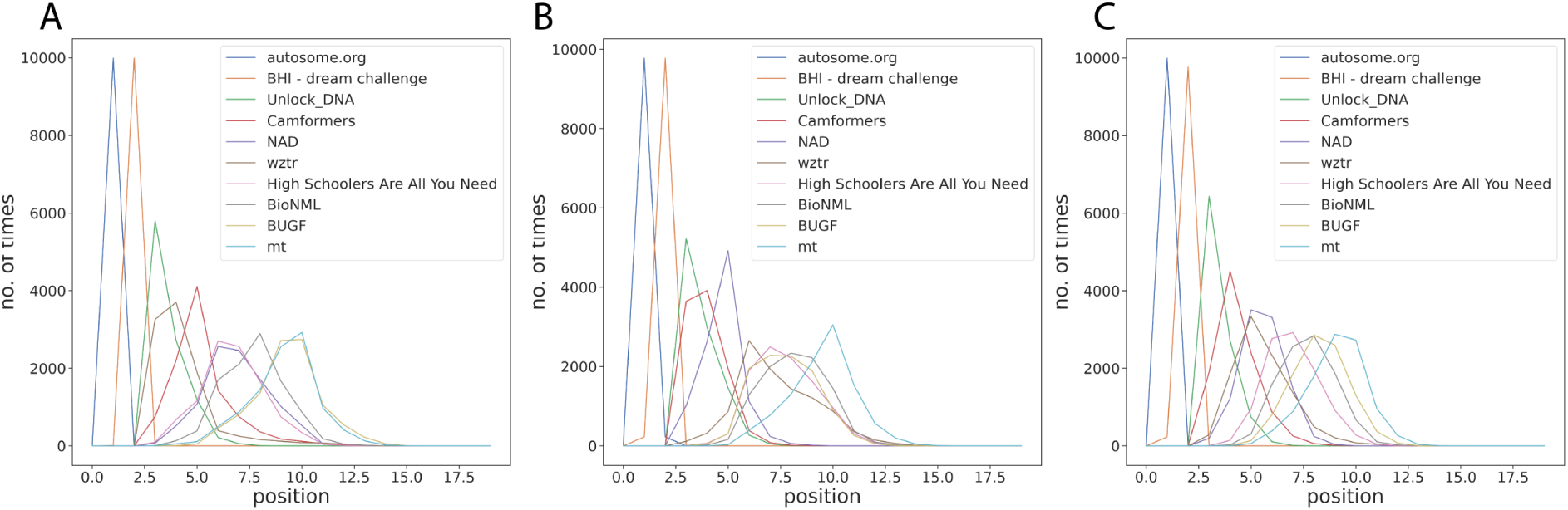
Bootstrapping provides a robust comparison of the model predictions. **(A,B,C)** Frequency (y-axes) of ranks (x-axes) in (A) Pearson Score, (B) Spearman Score and combined frequency (sum of Pearson Score rank and Spearman Score rank) of ranks for *n*=10,000 samples from the test dataset for the top-performing teams.

**Supplementary Figure 3:**
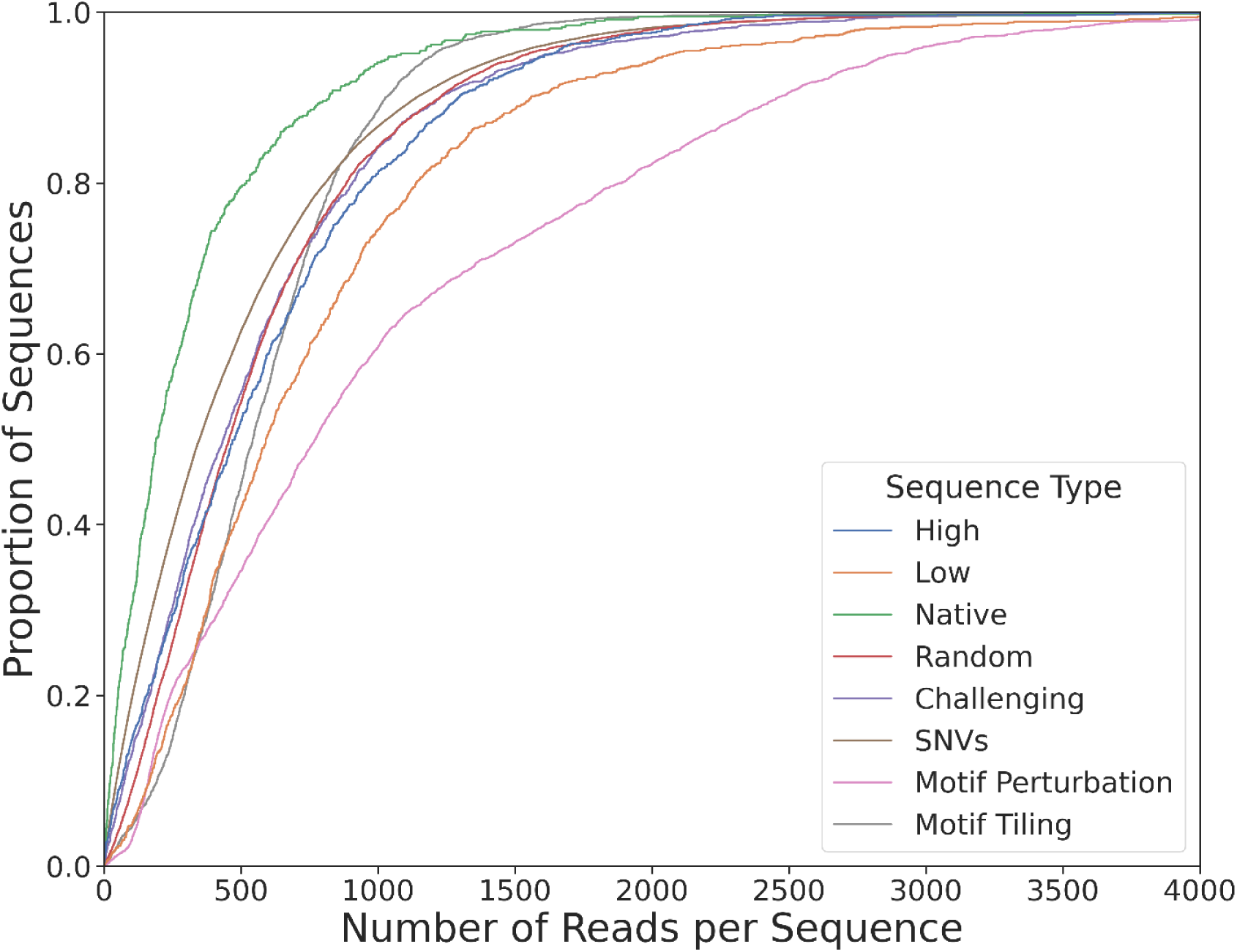
Library coverage differs between sequence subsets and is lowest for native sequences. Cumulative proportion (*y*-axis) of the number of reads per sequence (*x*-axis) for different sequence types (colours).

**Supplementary Figure 4:**
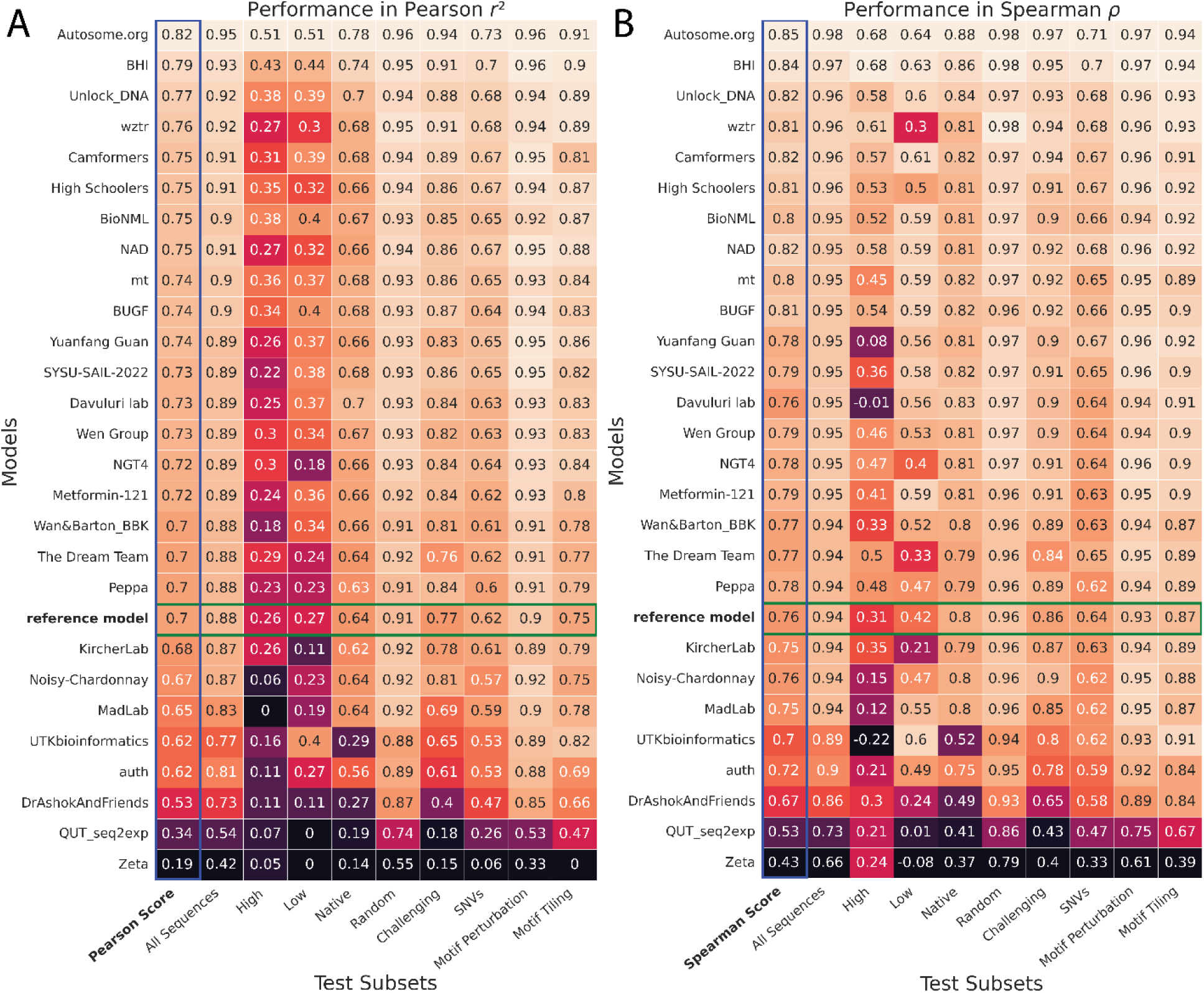
Performance of the teams in each test data subset. (A,B) Model performance (colour and numerical values) of each team (y-axes) in each test subset (x-axes), for (A) Pearson *r*^2^ and (B) Spearman ⍴. Heatmap colour palettes are min-max normalized column-wise.

**Supplementary Figure 5:**
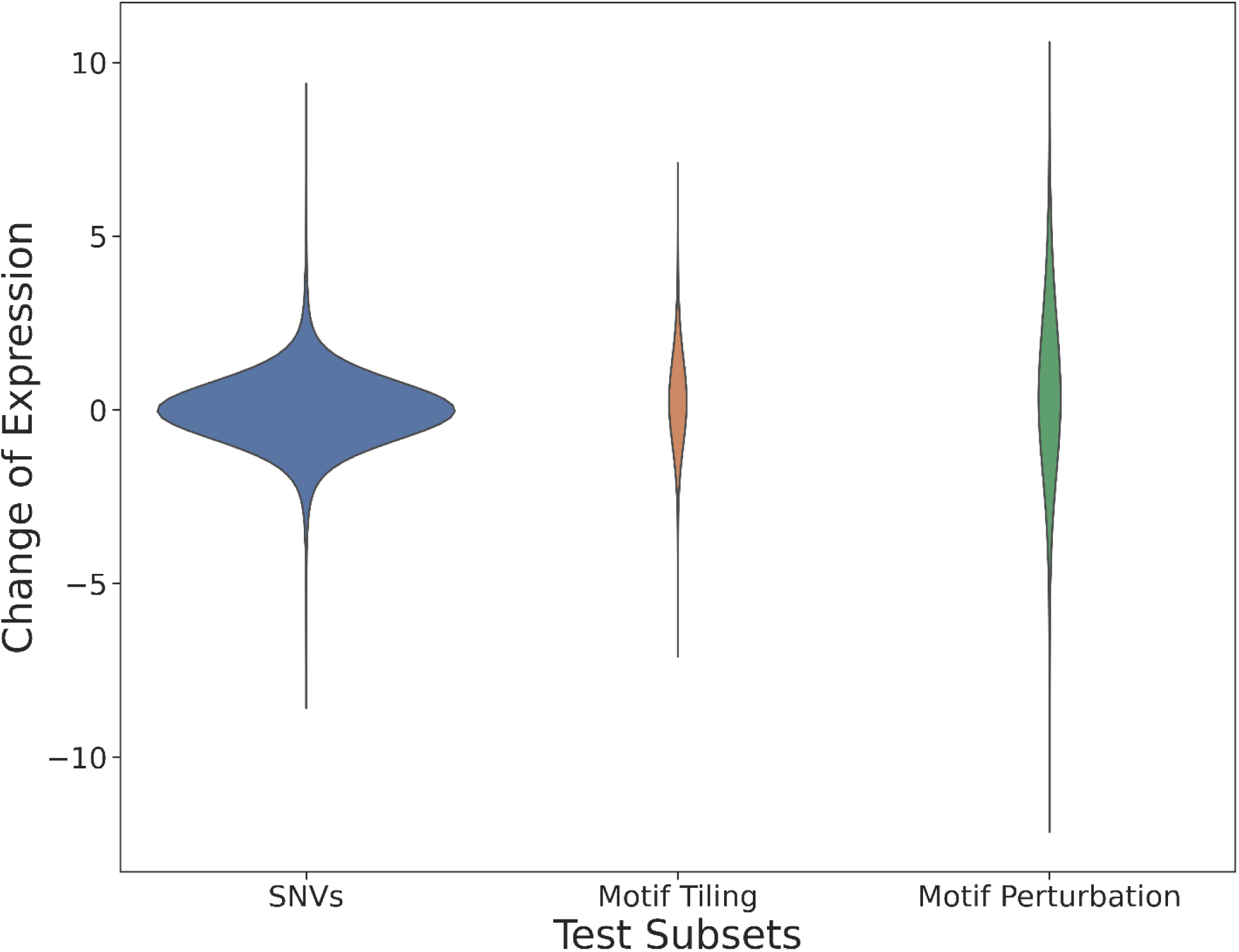
Expression changes (*y*-axis) are biggest for motif perturbation, smallest for SNVs, and intermediate for motif tiling.

**Supplementary Figure 6:**
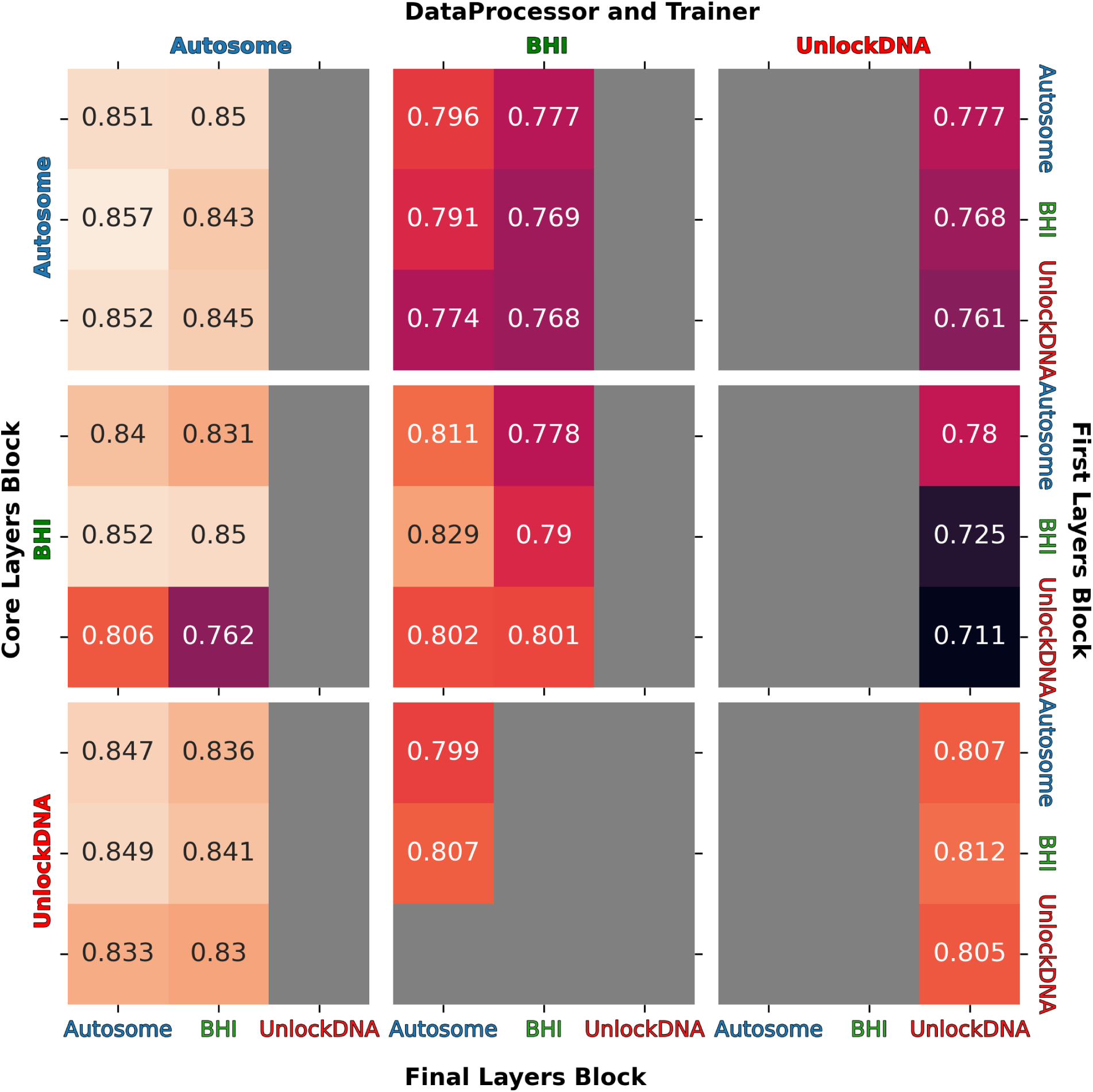
Performance in Spearman Score from the *Prix Fixe* runs for different possible combinations of the top three DREAM Challenge models. Modules are indicated on the axes, with Data Processor and Trainer models on the top *x* axis, Final Layer Block on the bottom *x* axis, Core Layers Block on the left *y* axis, and First Layers Block on the right *y* axis. Incompatible combinations and combinations that did not converge during training have been greyed out.

**Supplementary Figure 7:**
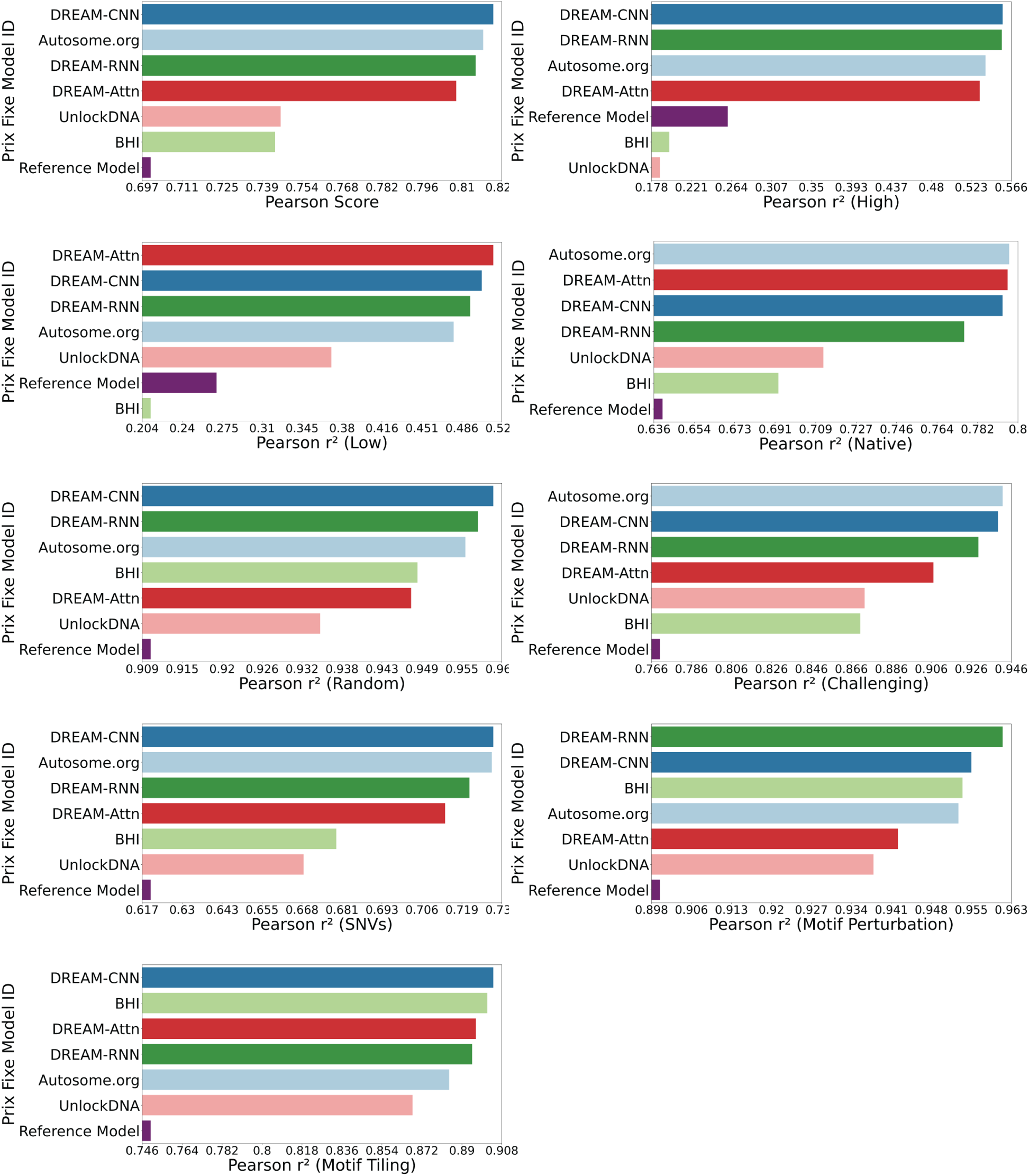
Performance (*x*-axes) of the top three DREAM Challenge models (*y*-axes) Autosome.org, BHI, and UnlockDNA-along with their best-performing counterparts (based on Core Layers Block) for different test subsets.

**Supplementary Figure 8:**
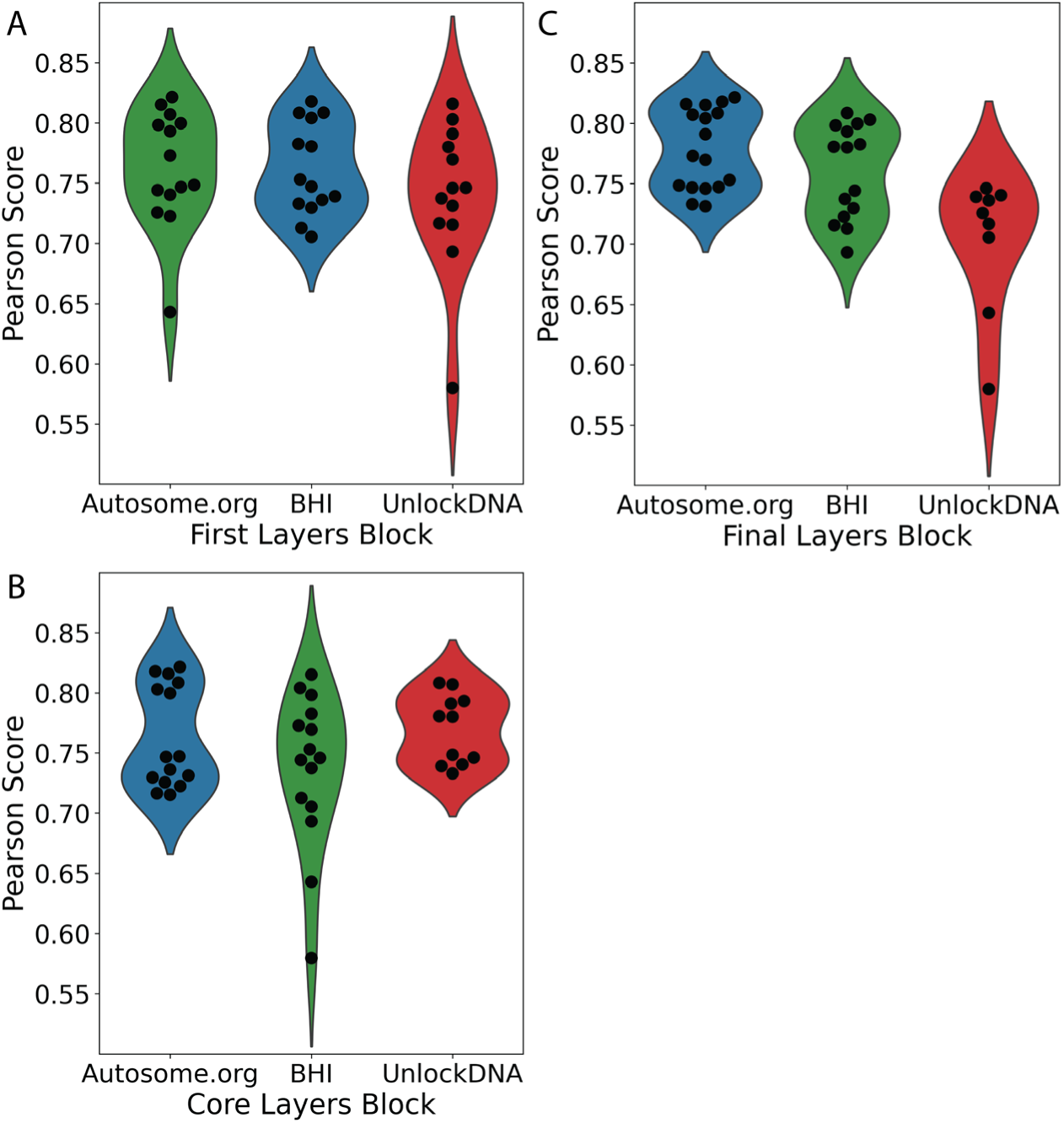
No clear winners in other module blocks. Performance of the different modules in Pearson Score (*y*-axis) for the different **(A)** First Layer, **(B)** Core Layers, and **(C)** Final Layers modules (*x*-axes and colours).

**Supplementary Figure 9:**
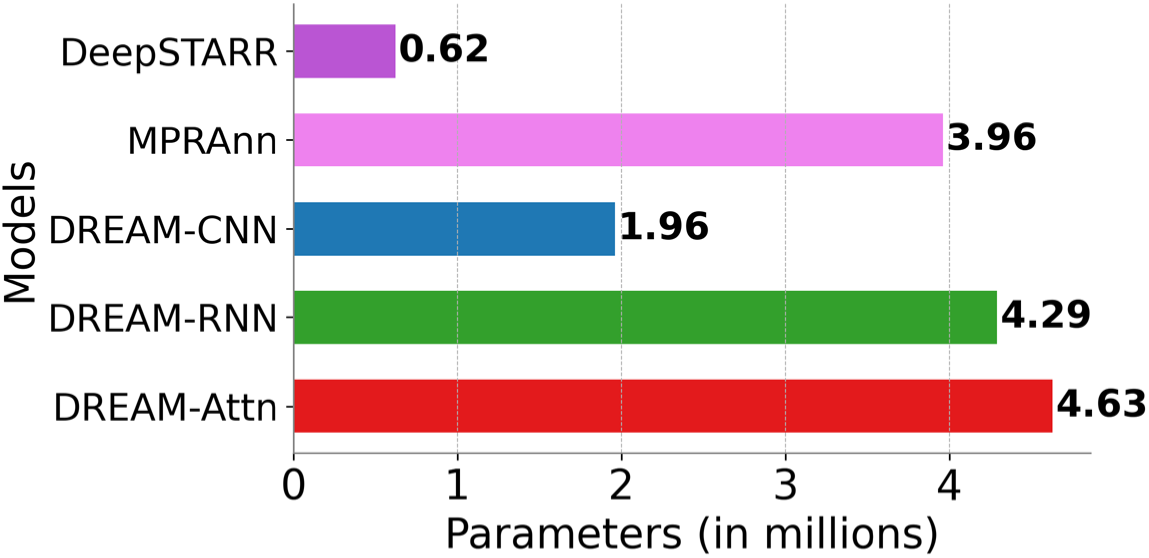
DREAM Optimized models have similar or fewer parameters than MPRAnn. Comparison of number of parameters (*x*-axis) for different models (*y*-axis) used in human MPRA prediction task (*51*).

## Acknowledgements

We extend our sincere gratitude to Jonathan Caton from Google Brain for his invaluable technical assistance in setting up the cloud resources for the challenge participants. Without his expert guidance and support, the successful organization of the challenge would not have been possible. We are also deeply grateful to TPU Research Cloud for providing the necessary cloud resources, which allowed us to ensure a level playing field for all challenge participants. We also thank Deep Genomics for providing travel grants to the top-performing teams in the challenge.

## Funding

This research was enabled in part by support provided by Google TRC, Digital Research Alliance of Canada, and Advanced Research Computing at the University of British Columbia. This research was supported by the Natural Sciences and Engineering Research Council of Canada (RGPIN-2020-05425 to CGD), the Stem Cell Network (ECR-C4R1-7 to CGD), and the Canadian Institute for Health Research (PJT-180537 to CGD). CGD is a Michael Smith Health Research BC Scholar and was supported by the NIH (grant no. K99-HG009920-01). AMR was supported by 4YF from the University of British Columbia. I.V.K. and D.P. were supported by RSF 20-74-10075. S.K. was supported by Institute of Information & communications Technology Planning & Evaluation (IITP) grant funded by the Korea government(MSIT) [NO.2021-0-01343, Artificial Intelligence Graduate School Program (Seoul National University)]. I.Y.K. was supported by National Research Foundation of Korea (NRF) grant funded by the Ministry of Science and ICT (RS-2023-00208284).

## Competing interests

E.D.V is the founder of Sequome, Inc. A.R. is an employee of Genentech and has equity in Roche. A.R. is a co-founder and equity holder of Celsius Therapeutics, an equity holder in Immunitas and, until 31 July 2020, was a scientific advisory board member of Thermo Fisher Scientific, Syros Pharmaceuticals, Neogene Therapeutics and Asimov. A.R. was an Investigator of the Howard Hughes Medical Institute when this work was initiated. The remaining authors declare no competing interests.

## Consortia

Random Promoter Dream Challenge Consortium members (in addition to the direct authors listed on the paper):

Susanne Bornelöv^1^, Fredrik Svensson^2^, Maria-Anna Trapotsi^1^, Duc Tran^3^, Tin Nguyen^3^, Xinming Tu^4^, Wuwei Zhang^4^, Wei Qiu^4^, Rohan Ghotra^5,6^, Yiyang Yu^5,6^, Ethan Labelson^6,7^, Aayush Prakash^8^, Ashwin Narayanan^9^, Peter Koo^6^, Xiaoting Chen^10^, David T. Jones^2^, Michele Tinti^11^, Yuanfang Guan^12^, Maolin Ding^34^, Ken Chen^34^, Yuedong Yang^34^, Ke Ding^13^, Gunjan Dixit^13^, Jiayu Wen^13^, Zhihan Zhou^14^, Pratik Dutta^15^, Rekha Sathian^15^, Pallavi Surana^15^, Yanrong Ji^14^, Han Liu^14^, Ramana V Davuluri^15^, Yu Hiratsuka^16^, Mao Takatsu^16^, Tsai-Min Chen^17,18^, Chih-Han Huang^19^, Hsuan-Kai Wang, Edward S.C. Shih^18^, Sz-Hau Chen^20^, Chih-Hsun Wu^21^, Jhih-Yu Chen^17^, Kuei-Lin Huang^22^, Ibrahim Alsaggaf^23^, Patrick Greaves^23^, Carl Barton^23^, Cen Wan^23^, Nicholas Abad^24^, Cindy Körner^24^, Lars Feuerbach^24^, Yichao Li^25^, Sebastian Röner^26^, Pyaree Mohan Dash^26^, Max Schubach^26^, Onuralp Soylemez^27^, Andreas Møller^28^, Gabija Kavaliauskaite^28^, Jesper Madsen^28^, Zhixiu Lu^29^, Owen Queen^29^, Ashley Babjac^29^, Scott Emrich^29^, Konstantinos Kardamiliotis^30^, Konstantinos Kyriakidis^30^, Andigoni Malousi^30^, Ashok Palaniappan^31^, Krishnakant Gupta^31^, Prasanna Kumar S^31^, Jake Bradford^32^, Dimitri Perrin^32^, Robert Salomone^32^, Carl Schmitz^32^, Chen JiaXing^33^, Wang JingZhe^33^, Yang AiWei^33^

Affiliations:

^1^ University of Cambridge

^2^ University College London

^3^ University of Nevada Reno

^4^ University of Washington

^5^ Syosset High School

^6^ Cold Spring Harbor Laboratory

^7^ Friends Academy

^8^ Half Hollow Hills High School

^9^ Jericho High School

^10^ Cincinnati Children’s Hospital Medical Center

^11^ The Wellcome Centre for Anti-Infectives Research, Dundee University

^12^ University of Michigan

^13^ Australia National University

^14^ Northwestern University

^15^ Stony Brook University

^16^ Niigata University School of Medicine

^17^ National Taiwan University

^18^ Academia Sinica

^19^ ANIWARE

^20^ Development Center for Biotechnology, Taipei

^21^ National Chengchi University

^22^ School of Medicine, China Medical University

^23^ Birkbeck, Univerity of London

^24^ German Cancer Research Center, Heidelberg

^25^ St. Jude Children’s Research Hospital

^26^ Berlin Institute of Health at Charité – Universitätsmedizin Berlin

^27^ Global Blood Therapeutics

^28^ University of Southern Denmark

^29^ University of Tennessee at Knoxville

^30^ Aristotle University of Thessaloniki

^31^ SASTRA University

^32^ Queensland University of Technology

^33^ Beijing Normal University-Hong Kong Baptist University United International College

^34^ Sun Yat-sen University, China

## Notes

https://zenodo.org/records/10633252

https://www.synapse.org/#!Synapse:syn28469146/wiki/

https://github.com/de-Boer-Lab/random-promoter-dream-challenge-2022

### Summary of Updates

All top-performing models from the challenge used neural networks, but diverged in architectures and novel training strategies, tailored to genomics sequence data. To dissect how architectural and training choices impact performance, we developed the Prix Fixe framework to divide any given model into logically equivalent building blocks. We tested all possible combinations for the top three models and observed performance improvements for each. The DREAM Challenge models not only achieved state-of-the-art results on our comprehensive yeast dataset but also consistently surpassed existing benchmarks on Drosophila and human genomic datasets. Overall, we demonstrate that high-quality gold-standard genomics datasets can drive significant progress in model development.

